# Choosing subsamples for sequencing studies by minimizing the average distance to the closest leaf

**DOI:** 10.1101/017822

**Authors:** Jonathan T. L. Kang, Peng Zhang, Sebastian Zöllner, Noah A. Rosenberg

**Affiliations:** Department of Biology, Stanford University, Stanford, CA 94305; Center for Inherited Disease Research, Johns Hopkins University, Baltimore, MD 21224; Department of Biostatistics, and Department of Psychiatry, University of Michigan, Ann Arbor, MI 48109

## Abstract

Imputation of genotypes in a study sample can make use of sequenced or densely genotyped external reference panels consisting of individuals that are not from the study sample. It can also employ internal reference panels, incorporating a subset of individuals from the study sample itself. Internal panels offer an advantage over external panels, as they can reduce imputation errors arising from genetic dissimilarity between a population of interest and a second, distinct population from which the external reference panel has been constructed. As the cost of next-generation sequencing decreases, internal reference panel selection is becoming increasingly feasible. However, it is not clear how best to select individuals to include in such panels. We introduce a new method for selecting an internal reference panel—minimizing the average distance to the closest leaf (ADCL)—and compare its performance relative to an earlier algorithm: maximizing phylogenetic diversity (PD). Employing both simulated data and sequences from the 1000 Genomes Project, we show that ADCL provides a significant improvement in imputation accuracy, especially for imputation of sites with low-frequency alleles. This improvement in imputation accuracy is robust to changes in reference panel size, marker density, and length of the imputation target region.

## INTRODUCTION

Owing to the existence of genetic variation within species, geneticists routinely make choices about which individuals, inbred strains, or representatives of populations or breeds merit prioritization for genotyping or DNA sequencing. Often, such choices, though typically made by informal criteria, reflect an explicit or implicit goal of maximizing the potential for extrapolating the information in the genotyped or sequenced individuals to all members of a breed, population, or species of interest.

Genotype imputation algorithms infer unobserved genotypes by matching a set of markers to the haplotype patterns observed in a reference sample (LI *et al*. 2009; MARCHINI AND HOWIE 2010), adding a new dimension to these choices. Reference panels that are used to facilitate genotype imputation in other individuals beyond the members of the panels themselves can often be optimally selected to formally maximize the imputed genotypic information obtained about those other individuals of interest (KANG AND MARJORAM 2012; ZHANG *et al*. 2013; PEIL *et al*. 2015). The evaluation of alternative ways to select imputation reference panels thus provides an approach for making sample choices for major genotyping or sequencing studies more systematically generalizable.

When conducting genotype imputation studies in a population sample, reference panels have generally been selected from databases external to the sample, such as the 1000 Genomes Project (1000 GENOMES PROJECT CONSORTIUM 2010) and the International HapMap Consortium (INTERNATIONAL HAPMAP CONSORTIUM 2005) databases. As a result of the rapidly decreasing cost of sequencing, however, it has become increasingly possible to carry out *internal* reference panel selection, in which additional sequencing is performed on a subset of the study sample, and the sequenced subset is then used to impute the remaining haplotypes. The use of reference sequences that originate from the study sample itself can reduce the potential mismatch of ancestral backgrounds between sample and reference populations, decreasing imputation errors. It also allows for genetic variants unique to the sample population to be successfully imputed (FRIDLEY *et al*. 2010; ZHANG *et al*. 2013).

Previous studies have observed that a mismatch in population origins between reference panels and study samples can reduce imputation accuracy compared to when they originate from the same or similar populations (HUANG *et al*. 2009, 2011; LI *et al*. 2010; PAŞANIUC *et al*. 2010; SHRINER *et al*. 2010; SURAKKA *et al*. 2010). JEWETT *et al*. (2012) demonstrated, using a coalescent model, that with other variables held constant, smaller internal reference panels are often likely to outperform larger external reference panels, despite the difference in panel size. Empirical studies have also shown that using an internal reference panel drawn from a subset of the sample under study, in addition to an external reference panel, gives rise to an increase in imputation accuracy over just using the external reference panel alone (FRIDLEY *et al*. 2010; SAMPSON *et al*. 2012; KREINER-MØLLER *et al*. 2015).

The value of internal reference panels for imputation studies raises the question of how an internal panel should be selected. Two recent studies have proposed maximizing “phylogenetic diversity” (PD) as a criterion for internal reference panel selection (KANG AND MARJORAM 2012; ZHANG *et al*. 2013). In this approach, the phylogenetic diversity of a set of haplotypes is defined as the total branch length of a tree spanned by the haplotypes (FAITH 1992; HARTMANN AND STEEL 2007). Given a panel size, the goal is to select the subset of haplotypes whose subtree yields the longest total branch length. Conceptually, the idea of seeking a maximally diverse subset of haplotypes in the reference panel aims to sample haplotypes that best cover the full range of haplotypes observed in the sample. The maximum-PD panel, by choosing haplotypes from different regions of the tree of haplotypes (FIGURE 1B), is more likely than a random panel to supply the necessary diversity to impute sites localized in a subgroup within the entire sample population. ZHANG *et al*. (2013) showed, using simulated sequence data and data from the 1000 Genomes Project, that by using the maximum-PD panel, higher imputation accuracy is obtained, and more sites are imputed as polymorphic in the sample population, than if the reference panel consists of randomly-selected haplotypes.

**FIGURE 1:**
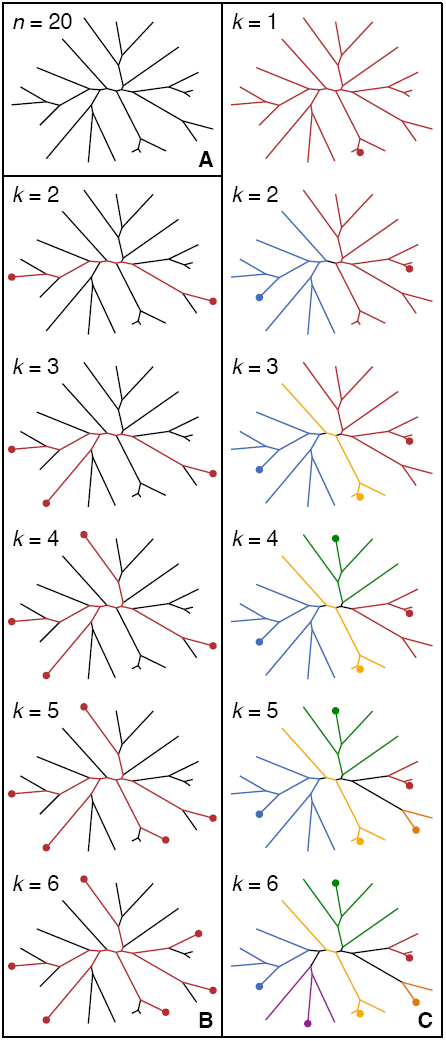
Reference panels for an example tree with *n* = 20 haplotypes. **(A)** An example tree. **(B)** The maximum-PD panel. **(C)** The minimum-ADCL panel. In **(B)** and **(C)**, the haplotypes selected for a given panel size *k* are represented by a dot at the tips. In **(C)**, each selected haplotype is assigned a color, and all other branches share a color with the closest selected haplotype.

Despite the utility of maximizing PD as a method for the selection of an internal reference panel, other approaches focusing on different principles might be preferable. Because the algorithm explicitly chooses haplotypes that are genetically distant from one another, long, pendant branches of the tree, if present, are likely to be chosen (BORDEWICH *et al*. 2008). The haplotypes associated with such branches might not be representative of the sample at large. These haplotypes might contain a large amount of sequencing error or missing data, and their inclusion in the reference panel might not contribute substantially to an increase in imputation accuracy. Even if they have high-quality data, such haplotypes are relatively unique in the sample, and therefore might assist as imputation templates only for a small number of sampled lineages.

PD can be viewed as emphasizing “diversity” of the internal reference panel rather than “representativeness.” To determine if an alternative focused on identifying the most representative subsample for use as the internal reference panel is preferable, we explore a new method: minimizing the average distance to the closest leaf (ADCL), which identifies reference haplotypes based on their genetic proximity to the rest of the sample haplotypes. We compare the imputation accuracy of the maximum-PD, minimum-ADCL, and random reference panels on both simulated data and data from the 1000 Genomes Project, and find that the minimum-ADCL panel consistently provides higher imputation accuracy, irrespective of changes to parameters such as reference panel size, marker density, and sequence length.

## METHODS

### Maximizing phylogenetic diversity (PD)

Given a tree of *n* haplotypes, to select a reference panel of haplotypes whose subtree spans the longest branch length, ZHANG *et al*. (2013) considered a greedy algorithm that takes as inputs the tree and a parameter *k ≤ n*, the desired number of haplotypes for the panel. Let *X* be the *k*-element subset of the sample haplotypes chosen for the reference panel, and let *T*_*X*_ be the subtree spanned by the haplotypes in *X*. The algorithm first selects the haplotype pair that is phylogenetically most distant (i.e. largest pairwise branch length), and adds both haplotypes to *X*. *T*_*X*_ now consists of a single pair of branches. Sequentially, the haplotype that is the most distant from *T*_*X*_ is placed into *X*, updating *T*_*X*_ with each inclusion. This process continues until the required *k* haplotypes have been selected (FIGURE 1B).

PARDI AND GOLDMAN (2005) and STEEL (2005) proved that among all possible subsets of size *k ≤ n* haplotypes from the study sample, the greedy algorithm achieves the globally maximal PD. Thus, the selection of the “most diverse” reference panel is computationally efficient, as there is no need to exhaustively examine all possible panels of size *k* in order to arrive at the correct solution. In addition, because the selection algorithm is greedy, the haplotypes in the reference panel can be ranked by their order of inclusion, in which every haplotype added contributes a non-increasing amount of PD. The maximum-PD panels of size 2 to *k* form a series of nested sets, and all previously selected haplotypes in a panel of size smaller than *k* will also be included in a panel of size *k*.

### Minimizing the average distance to the closest leaf (ADCL)

#### Overview of ADCL

Instead of focusing on diversity in the selected set and targeting the potential for accurate imputation of unusual haplotypes, the minimum-ADCL algorithm focuses on representativeness, aiming to maximize imputation accuracy of typical haplotypes likely to appear in a sample. The problem can be viewed as choosing the haplotypes that are, on average, genealogically closest to the remaining haplotypes not included in the reference panel. As in the case of PD, the algorithm takes as inputs a tree of the *n* haplotypes in the study sample, and a parameter *k ≤ n*, indicating the desired reference panel size.

Let *H* be the set of *n* haplotypes, and let *X* be the selected *k*-element subset of *H*. The objective is then to find *X* such that the branch-length distance from a randomly-chosen haplotype in *H* to its closest neighboring haplotype in *X* is minimized over all possible *k*-element subsets of *H* (MATSEN *et al*. 2013). Note that because the haplotypes in *X* are also in *H*, each of these haplotypes is its own closest neighbor, and we can equivalently consider either *H* or *H \ X*. In essence, the goal is to return a set of reference panel haplotypes that occupy the most central positions within clusters of the tree (FIGURE 1C).

In a detailed study of ADCL, MATSEN *et al*. (2013) demonstrated that unlike when choosing the subset that maximizes PD, the greedy algorithm need not give rise to the globally-optimal ADCL solution. It is therefore necessary to produce alternative algorithms that seek to minimize ADCL. Note that because the greedy algorithm is not applicable, the haplotypes selected cannot be ranked by their order of inclusion, as a haplotype included in a subset of size smaller than *k* is not necessarily also included in a subset of size *k* (FIGURE 1C).

#### Adapted partitioning-around-medoids (PAM) algorithm for minimizing ADCL

MATSEN *et al*. (2013) described two algorithms which, for a given set of haplotypes, seek to produce the subset of size *k* that minimizes ADCL. The first approach leverages similarities between the problem of minimizing ADCL and the technique known as *k*-medoids clustering (KAUFMAN AND ROUSSEEUW 1987). In the *k*-medoids problem, a set of data points is partitioned into *k* clusters, where *k* is predetermined. Within each cluster, a single point is designated as the center. The *k*-medoids clustering method is similar to *k*-means clustering. In the *k*-medoids approach, however, each cluster center is chosen from the original set of data points, whereas *k*-means has no such restriction. The objective function to be minimized in the *k*-medoids problem is the distance from a random data point to the center of the cluster to which it is assigned. A cluster center can be viewed as the data point most representative of the remainder of the data points within the cluster.

It is then clear how the problem of minimizing ADCL is analogous to the *k*-medoids problem. A data point is a haplotype, and distances between data points are branch-length (patristic) distances between haplotypes. The *k* cluster centers are akin to the *k* haplotypes that are selected.

As with minimizing ADCL, there is no greedy algorithm that solves the *k*-medoids problem, and obtaining the globally optimal solution has been demonstrated to be NP-hard (SHENG AND LIU 2004). A widely-used *k*-medoids heuristic algorithm is the partitioning-around-medoids (PAM) algorithm (THEODORIDIS AND KOUTROUMBAS 2008), which works by randomly selecting *k* medoids from the original set of *n* data points, and then minimizing the objective function via hill-climbing. One iteration of the algorithm consists of looping over all *k*(*n* − *k*) possible pairs containing a medoid and non-medoid, exchanging the medoid statuses of the points in the pair, and recording the new value of the objective function from the updated arrangement. Among all *k*(*n − k*) proposed exchanges, the single exchange that leads to the lowest-cost configuration is chosen. The algorithm then enters a new iteration, and the process repeats until no further changes to the set of medoids take place.

The first approach MATSEN *et al*. (2013) considered for minimizing ADCL is an adaptation of the PAM algorithm. First, the set *X* of haplotypes included in the reference panel is initialized by randomly selecting, without replacement, *k* haplotypes from the initial set *H* of *n* haplotypes. Next, the following loop over the haplotypes *x*_1_,…,*x*_*k*_ ∈ *X* is executed until no exchanges occur for one complete iteration over every *x*_*i*_ ∈ *X*:

1. For a haplotype *x*_*i*_ ∈ *X*, remove it from *X* and attempt to replace it with every other *y ∈ H \ X* in its place.
2. Keep the best such exchange if it decreases ADCL.
3. Continue with *x*_*i*+1_ ∈ *X*. In the case of *x*_*k*_, continue with *x*_1_.

This method for minimizing ADCL differs from the original formulation of the PAM algorithm in that it evaluates potential exchanges one medoid at a time, instead of examining all *k*(*n* − *k*) medoid/non-medoid pairs before finding the exchange that most decreases the objective function (MATSEN *et al*. 2013). Because each step in the iteration causes the value of ADCL to either stay constant or decrease, the solution is guaranteed to converge on a local minimum. However, the algorithm remains a heuristic approach, and the minimum-ADCL solution it achieves could depend on the specific haplotypes selected during random initialization. Hence, the global minimum might not always be found.

Alongside the adapted PAM algorithm, MATSEN *et al*. (2013) also developed a second approach: an exact but more computationally-intensive algorithm that is guaranteed to find the global-minimum ADCL solution. Both algorithms were implemented in the `rppr` binary in the `pplacer` suite of programs. Comparing between the two, MATSEN *et al*. (2013) demonstrated that for their simulated test sets, the adapted PAM algorithm only rarely gets trapped in local minima. For computational efficiency, we therefore chose to use the adapted PAM algorithm rather than the slower exact algorithm, first testing that in our setting, multiple runs of the adapted PAM algorithm with different initial seeds select a large percentage of the same haplotypes (see RESULTS).

### Simulated sequence data

To evaluate how the maximum-PD and minimum-ADCL panels perform relative to one another, we analyzed simulated data sets produced by the coalescent-based sequence sampling program `ms` (HUDSON 2002), closely following the parameters used by ZHANG *et al*. (2013) to ensure that the results are comparable.

First, we independently generated 50 data sets, each consisting of 2000 1Mb haplotypes, assuming a constant effective population size of *N*_*e*_ = 10, 000, a mutation rate of *µ* = 10^−8^ per site per generation, and a recombination rate of *ρ* = 10^−8^ per site per generation. The parameter values provided to `ms` were as follows: `nsam` = 2000, `nreps` = 50, `-t` = 400, `-r` = 400 and `nsites` = 10^6^. From the simulated data sets, we removed all singleton sites to ensure that the sequence data were truly imputable. Within a data set, if the *n* = 2000 haplotypes contained *q* polymorphic sites after excluding the singletons, we randomly selected, without replacement, *s* = 300 of the *q* sites, each with minor allele frequency (MAF) greater than 0.1. These markers were treated as genotyped. The remaining *q* – *s* sites were masked.

Following ZHANG *et al*. (2013), we calculated the pairwise Hamming distances between the *n* = 2000 haplotypes in each of the 50 data sets, based on the genotype information at only the *s* = 300 randomly-selected markers. With these distances, we then used the software `rapidnj` (SIMONSEN *et al*. 2008) to construct a neighbor-joining tree (SAITOU AND NEI 1987) of the haplotypes. Note that it was possible, as a result of random sampling, for two or more haplotypes to be identical at all *s* markers. In such a case, a leaf in the tree would represent more than one haplotype.

Using the python library `dendropy` (SUKUMARAN AND HOLDER 2010), we calculated the patristic distance matrix for each neighbor-joining tree. We then applied the greedy algorithm to select the reference panel of size *k* = 200 that maximizes PD. Furthermore, on each neighbor-joining tree, we used the `rppr` binary in `pplacer` (MATSEN *et al*. 2013) to execute the adapted PAM algorithm, returning a reference panel of size *k* = 200 that minimizes ADCL. In cases for which either algorithm selected a leaf that represents more than one haplotype, one of the haplotypes was randomly chosen to be included in the panel.

In order to model diploid samples, we also created diploid reference panels for use with both the maximum-PD and minimum-ADCL algorithms. First, we randomly paired the *n* = 2000 haplotypes into 1000 diploid genomes. For the “diploid PD” panel, following ZHANG *et al*. (2013), we included diploid individuals carrying at least one of the top-ranked haplotypes into the panel until we reached the desired panel size *k*. More specifically, we proceeded down the list of *k* haplotypes in the maximum-PD panel, ranked based on the order of inclusion. At each step, we selected both the top-ranked haplotype and the haplotype with which it was paired (and which was not necessarily top-ranked) for the diploid panel, if they had not already been picked previously. We continued this process until *k/*2 diploid genomes were selected, for a total of *k* haplotypes.

Unlike in maximum-PD panels, haplotype sets in minimum-ADCL panels are not nested. Therefore, we cannot use the same process to construct the “diploid ADCL” panel. To address this problem, we first constructed, again using `rppr`, a half-sized minimum-ADCL panel of size *k*/2 = 100. Each haplotype in the half-sized panel, along with the haplotype with which it was paired, was then included in the diploid panel. In the event that both haplotypes of a diploid genome were in the half-sized panel, they were each only chosen once. If the diploid panel was not fully filled at the end of this process, then haplotype pairs were randomly taken from the previously unselected diploid genomes until the requisite panel size of *k*/2 diploid genomes was reached.

For comparison, for each of the 50 data sets, we also generated 1000 random reference panels by sampling, without replacement, *k* = 200 of the original *n* = 2000 haplotypes, giving a total of 1004 reference panels. A diagram of the simulation pipeline appears in FIGURE 2.

**FIGURE 2:**
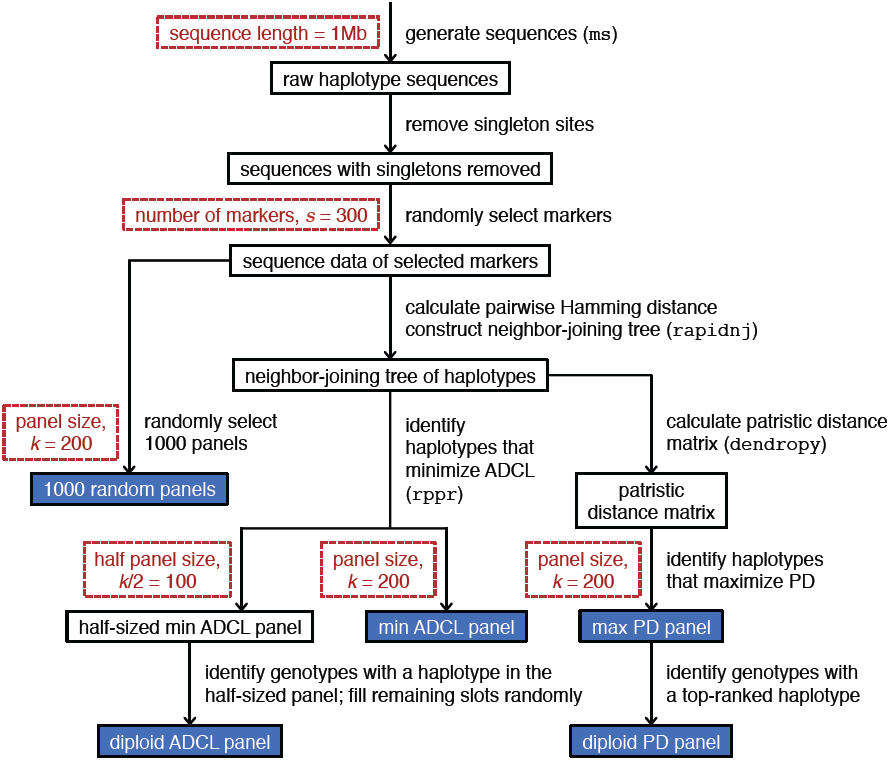
A schematic diagram of the pipeline used to generate the simulated data. The red boxes each represent a parameter choice, and the blue boxes represent the 1004 reference panels used in our evaluation.

For each of the *k* haplotypes in a reference panel, we unmasked the genotypes at the *q* – *s* masked sites and used the resulting full sequences as a reference to perform imputation, under the assumption that the haplotypes represent sequences with resolved phasing. Following ZHANG *et al*. (2013), to avoid edge effects and to improve imputation accuracy, within each 1Mb haplotype, we imputed only the middle 100kb segment, while still retaining the markers in both 450kb flanking regions (LI *et al*. 2010). Similar to ZHANG *et al*. (2013), we used the program `minimac` (HOWIE *et al*. 2012) to perform imputation. The parameter values entered into `minimac` were as follows: `--rounds` = 5 and `--states` = 200.

For each choice of reference panel, we evaluated imputation accuracy at the *r* imputed sites (masked sites within the middle 100kb segment) over the *n*/2 diploid genomes, applying a discordance metric. At imputed site *j* in diploid genome *i*, we define *g*_*ij*_ and *ĝ*_*ij*_ to be the true and imputed genotypes respectively. Both *g*_*ij*_ and *ĝ*_*ij*_ take on values in {0, 1, 2}, corresponding to the number of copies of an arbitrarily chosen allele at that specific site. The discordance rate *D* across all sites is given by

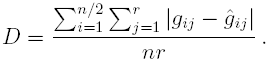

We also compute the discordance rate *H* across all true heterozygous genotypes (*g*_*ij*_ = 1):

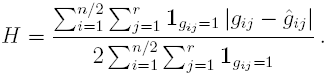

In addition, based on the MAF values of their constituent alleles, as computed in the full set of 2000 haplotypes, we further split the true heterozygous sites into three mutually exclusive MAF bins: 0 < MAF < 0.1 (low), 0.1 ≤ MAF < 0.2 (medium), and 0.2 ≤ MAF ≤ 0.5 (high). This separation was performed in order to evaluate how the PD and ADCL algorithms perform across the spectrum of rare to common variants. Note also that the calculations of *D* and *H* sum over all *n*/2 diploid genomes, irrespective whether they have one, both, or neither of their haplotypes represented in the reference panel.

### 1000 Genomes Project sequence data

We also applied both the PD and ADCL algorithms to sequence data from the 1000 Genomes Project, available at http://csg.sph.umich.edu/abecasis/MACH/download/ 1000G-PhaseI-Interim.html. Following ZHANG *et al*. (2013), we considered *n* = 762 phased haplotypes from 381 diploid individuals with European ancestry: 87 Utah residents with Northern and Western European ancestry, 93 Finnish from Finland, 89 British from England and Scotland, 14 Iberians from Spain, and 98 Toscani from Italy.

We first removed all singleton sites from the data, and we then selected 30 1Mb segments that were approximately evenly spaced across chromosome 20, avoiding the centromere, telomeres, and adjacent areas. Study samples were then created using a similar procedure to that employed for the simulated data. For each of the 30 segments, we randomly selected *s* = 400 markers with MAF *>* 0.1 in the full set of 762 haplotypes, and masked the genotypes of the remaining sites. We then chose *k* = 120 haplotypes to include in the maximum-PD and minimum-ADCL reference panels, as well as in 1000 randomly-generated panels. For each choice of reference panel used for each segment, we imputed the middle 100kb, retaining the markers in both 450kb flanking regions. We then evaluated *D* and *H* analogously to the experiments with the simulated data.

## RESULTS

### Stability of the adapted PAM algorithm

Before considering the actual imputation results produced by the different algorithms for reference panel selection, we empirically validated the stability of the adapted PAM algorithm in choosing the minimum-ADCL panel. Beyond the initial run for each of our 50 simulated data sets, we repeated the selection of the minimum-ADCL panel five additional times. For each repetition, we executed the adapted PAM algorithm with a different starting seed, and then determined the number of haplotypes that were shared by the minimum-ADCL panels from both the initial run and the run with the modified seed.

When comparing two panels of 200 reference haplotypes drawn from a set of 2000 sample haplotypes, let *m* be the number of haplotypes that are shared by both panels (0 *≤ m ≤* 200). For each of the five replicates, we calculated the mean value of *m* across the 50 data sets, comparing each replicate to the initial run. All five mean values of *m* were observed to be *∼*179 (TABLE 1); for comparison, the mean of the hypergeometric distribution describing the number of haplotypes shared between two panels of size 200 independently drawn from a pool of 2000 is 20, with standard deviation 4.03. Therefore, despite changing the specific haplotypes used in randomly initializing the adapted PAM algorithm, most haplotypes eventually chosen for inclusion in the minimum-ADCL panel remain the same. This result suggests that the adapted PAM algorithm is in fact stable, and in subsequent analysis, we consider only a single starting seed.

**TABLE 1:** Mean and standard deviation of the number of shared haplotypes across 50 data sets in each of five replicates For the five replicates, each with a different starting seed, we compared the minimum-ADCL panels from the initial run of the adapted PAM algorithm and the minimum-ADCL panels using the different seed. The table shows the mean (out of 200) and standard deviation of the number of shared haplotypes across the 50 data sets.

| Replicate | Mean | Standard deviation |
| --- | --- | --- |
| 1 | 179.40 | 4.1991 |
| 2 | 178.56 | 4.9494 |
| 3 | 179.38 | 4.5888 |
| 4 | 179.00 | 4.7208 |
| 5 | 178.58 | 4.8910 |

### Polymorphic sites in reference panels

For each of the 1004 reference panels, we evaluated the number of masked sites within the imputed 100kb segment that were polymorphic. This calculation is important because only sites that are polymorphic in the reference panel can produce a meaningful imputation result for the remainder of the study sample. Summing across all 50 data sets, we detected a total of 12,851 masked sites within the 100kb segment of interest. We then compared how many of those masked sites appear as polymorphic in the maximum-PD panel, the minimum-ADCL panel, and a single random panel.

Of the 12,851 masked sites, 8879 sites (69.09%) were polymorphic in all three reference-panel types. Of the 3972 remaining sites, 1138 (8.86%) were polymorphic in both the maximum-PD and minimum-ADCL panels, 244 (1.90%) were polymorphic in both the maximum-PD and random panels, and 374 (2.91%) were polymorphic in both the minimum-ADCL and random panels. In addition, 464 (3.61%), 473 (3.68%), and 391 (3.04%) sites were polymorphic in only the maximum-PD, minimum-ADCL, and random panels, respectively. Finally, 888 (6.91%) of the masked sites were monomorphic in all three panels (FIGURE 3).

**FIGURE 3:**
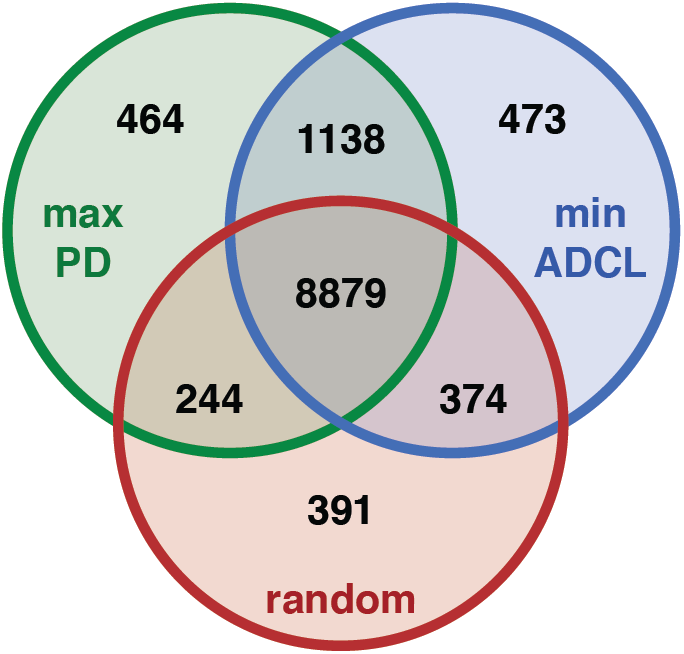
A Venn diagram showing the number of polymorphic sites returned by each panel type, out of a total of 12,851 masked sites. 888 sites were monomorphic in all three panels.

Overall, 10,725 sites (83.46%) were polymorphic in the 50 maximum-PD panels, 10,864 sites (84.54%) were polymorphic in the 50 minimum-ADCL panels, and 9888 sites (76.94%) were polymorphic in the 50 random panels. Using the two-tailed Wilcoxon signed-rank test, we found that both the maximum-PD and minimum-ADCL methods of panel selection identify substantially more polymorphic sites compared to choosing the reference panel randomly (*P* = 7.686 × 10^−10^ and *P* = 8.175 × 10^−10^, respectively).

### Polymorphic sites in imputed data sets

The maximum-PD and minimum-ADCL selection algorithms result in similar numbers of polymorphic sites as a fraction of the total number of masked sites in their respective reference panels. We next evaluated the number of imputed sites the two methods recovered as polymorphic. In each of the 50 simulated data sets, we calculated the percentage of masked sites that were polymorphic in the imputed sample, using the maximum-PD panel, the minimum-ADCL panel, the diploid PD panel, the diploid ADCL panel, and the same random panel used to assess the number of polymorphic sites within the reference panels.

FIGURE 4 compares the proportion of polymorphic sites imputed with combinations of the five reference panel types. In each panel of FIGURE 4, the random panel is used as a baseline for evaluating two of the other four panel selection methods.

**FIGURE 4:**
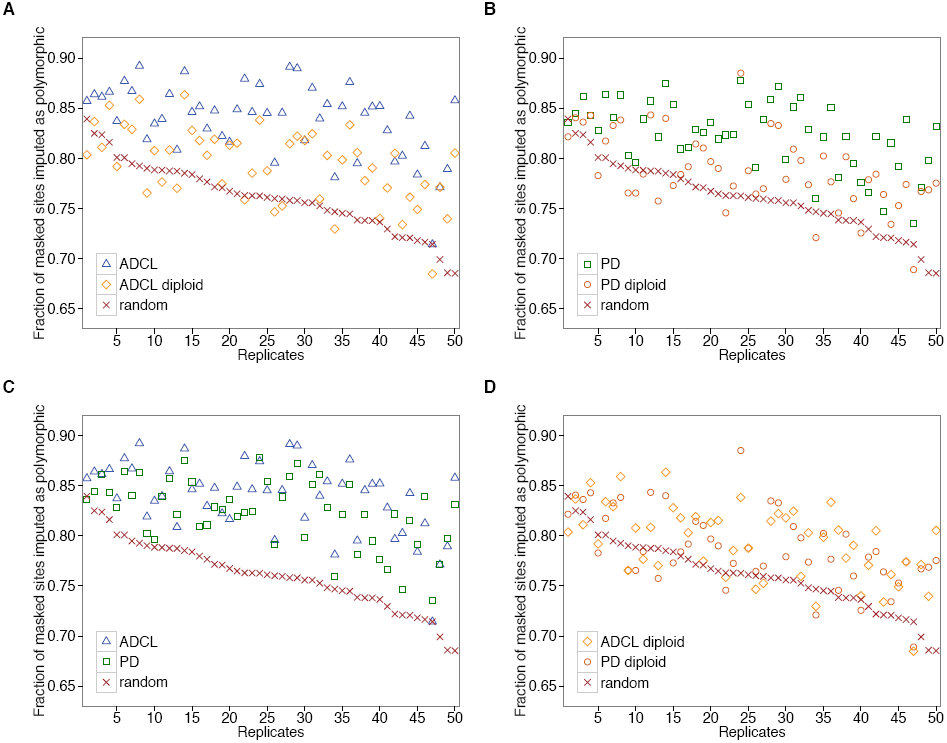
Fraction of masked sites imputed as polymorphic, using five different types of reference panels. Data are split into various graphs for ease of comparison. **(A)** ADCL versus ADCL diploid. **(B)** PD versus PD diploid. **(C)** ADCL versus PD. **(D)** ADCL diploid versus PD diploid. The 50 replicate data sets are sorted in decreasing order by the percentage of polymorphic sites recovered by imputations using the random reference panel.

We used the two-tailed Wilcoxon signed-rank test to evaluate differences in the fraction of sites identified as polymorphic by the different panel types. Both the maximum-PD and minimum-ADCL panels recover a significantly larger percentage of polymorphic sites compared with their respective diploid panels (*P* = 3.448 × 10^−9^ and *P* = 2.309 × 10^−9^, respectively). The minimum-ADCL panel also outperforms the maximum-PD panel (*P* = 4.944 × 10^−4^). However, the percentage of imputed sites that are polymorphic shows no significant difference when comparing the diploid PD and diploid ADCL panels (*P* = 0.1625).

### Discordance rates

As a measure of imputation accuracy, for each of the 50 simulated data sets, we separately calculated the discordance rate *D* across all sites that were imputed with the maximum-PD panel, the minimum-ADCL panel, the diploid PD panel, and the diploid ADCL panel. For a baseline, we also calculated the mean discordance rate over the 1000 randomly-selected reference panels. We are mainly interested in comparing the performance between the maximum-PD and minimum-ADCL panels, as well as between the diploid PD and diploid ADCL panels.

The discordance rates appear in FIGURE 5, and their mean values are summarized in TABLE 2. Again using the two-tailed Wilcoxon signed-rank test, the minimum-ADCL panel exhibits significantly lower discordance rates than the maximum-PD panel (*P* = 1.342 × 10^−9^). The diploid ADCL panel also has lower discordance rates than the diploid PD panel (*P* = 2.597 × 10^−3^). The minimum-ADCL, maximum-PD, diploid ADCL, and diploid PD panels all provide lower discordance rates than the mean of the 1000 randomly-selected panels (*P* = 7.789 × 10^−10^, 9.928 × 10^−10^, 8.797 × 10^−10^, and 4.920 × 10^−7^, respectively).

**TABLE 2:** Mean discordance rates between imputed and simulated genotypes, using the maximum (haploid) PD, minimum (haploid) ADCL, diploid PD, and diploid ADCL panels This table is obtained from the data in FIGURE 5. The comparison is performed over all sites, all heterozygous sites, and heterozygous sites falling into three different MAF groups. Also shown are the *P*-values of the two-tailed Wilcoxon signed-rank tests comparing the discordance rates of the PD and ADCL reference panels.

|  | Haploid panels |  |  | Diploid panels |  |  |
| --- | --- | --- | --- | --- | --- | --- |
| | PD (%) | ADCL (%) | $P$-value | PD (%) | ADCL (%) | $P$-value |
| All | 0.2003 | 0.1476 | $1.342 \times 10^{-9}$ | 0.2648 | 0.2304 | $2.597 \times 10^{-3}$ |
| Heterozygotes | 1.1295 | 0.7888 | $1.921 \times 10^{-9}$ | 1.4554 | 1.2523 | $1.360 \times 10^{-3}$ |
| MAF (0, 0.1) | 3.0374 | 2.1160 | $1.606 \times 10^{-9}$ | 3.8164 | 3.2103 | $1.871 \times 10^{-4}$ |
| MAF [0.1, 0.2) | 0.4363 | 0.3190 | $2.792 \times 10^{-3}$ | 0.6947 | 0.5582 | 0.0857 |
| MAF [0.2, 0.5] | 0.3518 | 0.2386 | $2.567 \times 10^{-5}$ | 0.4676 | 0.4335 | 0.4174 |

**FIGURE 5:**
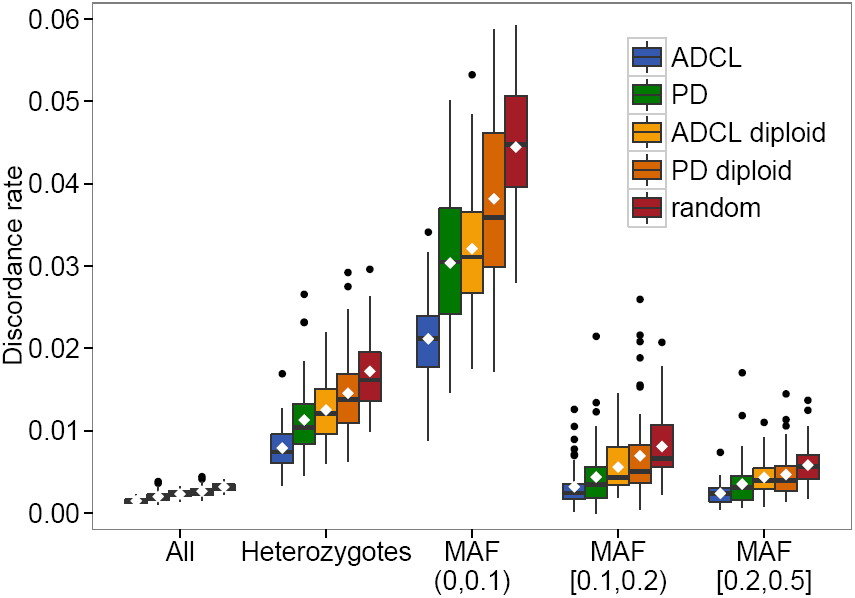
Box plots of discordance rates between imputed and simulated genotypes using the five different reference panel types. The mean discordance rate across the 50 replicates for each comparison group is indicated by a diamond, and the median discordance rate is indicated by a horizontal line. The *x*-axis separates the comparison over all sites, all heterozygous sites, and heterozygous sites falling into three different MAF groups.

To generate a discordance measure for low-frequency variants, we also calculated the discordance rate *H* across the heterozygous sites with 0 < MAF < 0.1. From FIGURE 5 and TABLE 2, we observe that the mean discordance rates are higher for low-MAF loci than they are for high-MAF loci. Nevertheless, compared to the maximum-PD panel, the minimum-ADCL panel still achieves significantly higher imputation accuracy on low-MAF heterozygotes (*P* = 1.606 × 10^−9^). The same relationship also holds between the diploid ADCL and diploid PD panels (*P* = 1.871 × 10^−4^). As was observed when considering all variants, the minimum-ADCL, maximum-PD, diploid ADCL, and diploid PD panels all have lower discordance rates than the mean of the 1000 random panels (*P* = 7.790 × 10^−10^, 1.264 × 10^−9^, 7.790 × 10^−10^, and 2.244 × 10^−6^, respectively).

### Discordance rates under different simulation settings

Following ZHANG *et al*. (2013), to investigate how different parameter choices might have affected the simulation results, we repeated the analysis taking into consideration (i) different reference panel sizes *k*, (ii) different marker densities *s*, and (iii) different target sequence lengths. When varying a parameter, we kept the other two parameters constant at their default values used in the initial analysis (reference panel size *k* = 200, number of markers per MB *s* = 300, imputation length = 100kb). The baseline for comparison here is the mean discordance rate over the 50 randomly-selected reference panels. Owing to runtime considerations, this number is smaller than the 1000 randomly-selected reference panels used to calculate the baseline mean discordance rate in the initial analysis. Box plots of the results are shown in FIGURE 6, and mean discordance rates of the various panel types over all sites and over the low-frequency variants appear in TABLES 3 and 4, respectively.

**TABLE 3:** Mean discordance rates between imputed and simulated genotypes for all sites, using the maximum (haploid) PD, minimum (haploid) ADCL, diploid PD, and diploid ADCL panels, under different input parameter choices The table is obtained from the data in FIGURES 6A, C and E. Also shown are the *P*-values of the two-tailed Wilcoxon signed-rank tests comparing the discordance rates of the PD and ADCL reference panels. The discordance rates and *P*-values from the initial analysis using *k* = 200, *s* = 300 and imputation length = 100kb are given in bold, with the values obtained from TABLE 2.

|  | Haploid panels |  |  | Diploid panels |  |  |
| --- | --- | --- | --- | --- | --- | --- |
| | PD (%) | ADCL (%) | $P$-value | PD (%) | ADCL (%) | $P$-value |
| Reference panel size, $k$ | | | | | | |
| $k = 100$ | 0.5150 | 0.3502 | $5.213 \times 10^{-9}$ | 0.6174 | 0.5284 | $2.257 \times 10^{-5}$ |
| **$k = 200$** | **0.2003** | **0.1476** | **$1.342 \times 10^{-9}$** | **0.2648** | **0.2304** | **$2.597 \times 10^{-3}$** |
| $k = 300$ | 0.0907 | 0.0811 | $2.767 \times 10^{-3}$ | 0.1501 | 0.1354 | 0.0167 |
| $k = 400$ | 0.0499 | 0.0498 | 0.7391 | 0.0924 | 0.0895 | 0.3417 |
| $k = 500$ | 0.0298 | 0.0312 | 0.2112 | 0.0584 | 0.0605 | 0.1765 |
| Number of markers per MB, $s$ | | | | | | |
| $s = 200$ | 0.2561 | 0.1810 | $1.378 \times 10^{-8}$ | 0.3409 | 0.2901 | $1.173 \times 10^{-4}$ |
| **$s = 300$** | **0.2003** | **0.1476** | **$1.342 \times 10^{-9}$** | **0.2648** | **0.2304** | **$2.597 \times 10^{-3}$** |
| $s = 400$ | 0.1647 | 0.1255 | $2.548 \times 10^{-8}$ | 0.2291 | 0.1946 | $4.176 \times 10^{-5}$ |
| $s = 500$ | 0.1529 | 0.1193 | $9.347 \times 10^{-10}$ | 0.2105 | 0.1863 | $7.284 \times 10^{-4}$ |
| $s = 600$ | 0.1503 | 0.1138 | $4.130 \times 10^{-9}$ | 0.1970 | 0.1746 | $6.975 \times 10^{-5}$ |
| Imputation length |  |  |  |  |  |  |
| **100kb** | **0.2003** | **0.1476** | **$1.342 \times 10^{-9}$** | **0.2648** | **0.2304** | **$2.597 \times 10^{-3}$** |
| 500kb | 0.2104 | 0.1494 | $7.789 \times 10^{-10}$ | 0.2650 | 0.2307 | $5.575 \times 10^{-6}$ |
| 1Mb | 0.2159 | 0.1538 | $7.790 \times 10^{-10}$ | 0.2755 | 0.2392 | $2.040 \times 10^{-8}$ |
| 2Mb | 0.2495 | 0.1653 | $7.789 \times 10^{-10}$ | 0.2914 | 0.2551 | $8.263 \times 10^{-9}$ |

**TABLE 4:** Mean discordance rates between imputed and simulated genotypes for all heterozygous sites with 0 < MAF < 0.1, using the maximum (haploid) PD, minimum (haploid) ADCL, diploid PD, and diploid ADCL panels, under different input parameter choices The table is obtained from the data in FIGURES 6B, D and F. Also shown are the *P*-values of the two-tailed Wilcoxon signed-rank tests comparing the discordance rates of the PD and ADCL reference panels. The discordance rates and *P*-values from the initial analysis using *k* = 200, *s* = 300 and imputation length = 100kb are given in bold, with the values obtained from TABLE 2.

|  | Haploid panels |  |  | Diploid panels |  |  |
| --- | --- | --- | --- | --- | --- | --- |
| | PD (%) | ADCL (%) | $P$-value | PD (%) | ADCL (%) | $P$-value |
| Reference panel size, $k$ | | | | | | |
| $k = 100$ | 7.2099 | 5.0056 | $3.358 \times 10^{-8}$ | 8.5331 | 7.3942 | $3.960 \times 10^{-4}$ |
| **$k = 200$** | **3.0374** | **2.1160** | **$1.606 \times 10^{-9}$** | **3.8164** | **3.2103** | **$1.871 \times 10^{-4}$** |
| $k = 300$ | 1.4783 | 1.2425 | $1.484 \times 10^{-4}$ | 2.1964 | 1.9086 | $3.817 \times 10^{-4}$ |
| $k = 400$ | 0.8394 | 0.8156 | 0.2972 | 1.3786 | 1.2816 | 0.0757 |
| $k = 500$ | 0.5494 | 0.5413 | 0.7502 | 0.9160 | 0.9010 | 0.7138 |
| Number of markers per MB, $s$ | | | | | | |
| $s = 200$ | 3.4597 | 2.3336 | $3.467 \times 10^{-9}$ | 4.3339 | 3.5993 | $8.581 \times 10^{-6}$ |
| **$s = 300$** | **3.0374** | **2.1160** | **$1.606 \times 10^{-9}$** | **3.8164** | **3.2103** | **$1.871 \times 10^{-4}$** |
| $s = 400$ | 2.6079 | 1.9485 | $3.270 \times 10^{-9}$ | 3.5185 | 2.9342 | $1.173 \times 10^{-5}$ |
| $s = 500$ | 2.4951 | 1.8300 | $1.810 \times 10^{-9}$ | 3.4146 | 2.8719 | $7.874 \times 10^{-5}$ |
| $s = 600$ | 2.4121 | 1.7891 | $7.368 \times 10^{-9}$ | 3.2507 | 2.7438 | $1.528 \times 10^{-5}$ |
| Imputation length |  |  |  |  |  |  |
| **100kb** | **3.0374** | **2.1160** | **$1.606 \times 10^{-9}$** | **3.8164** | **3.2103** | **$1.871 \times 10^{-4}$** |
| 500kb | 2.8998 | 2.0484 | $7.790 \times 10^{-10}$ | 3.5755 | 3.0995 | $1.605 \times 10^{-6}$ |
| 1Mb | 3.0180 | 2.1204 | $7.790 \times 10^{-10}$ | 3.7531 | 3.2439 | $1.231 \times 10^{-8}$ |
| 2Mb | 3.4652 | 2.2648 | $7.790 \times 10^{-10}$ | 3.9769 | 3.4637 | $9.806 \times 10^{-9}$ |

**FIGURE 6:**
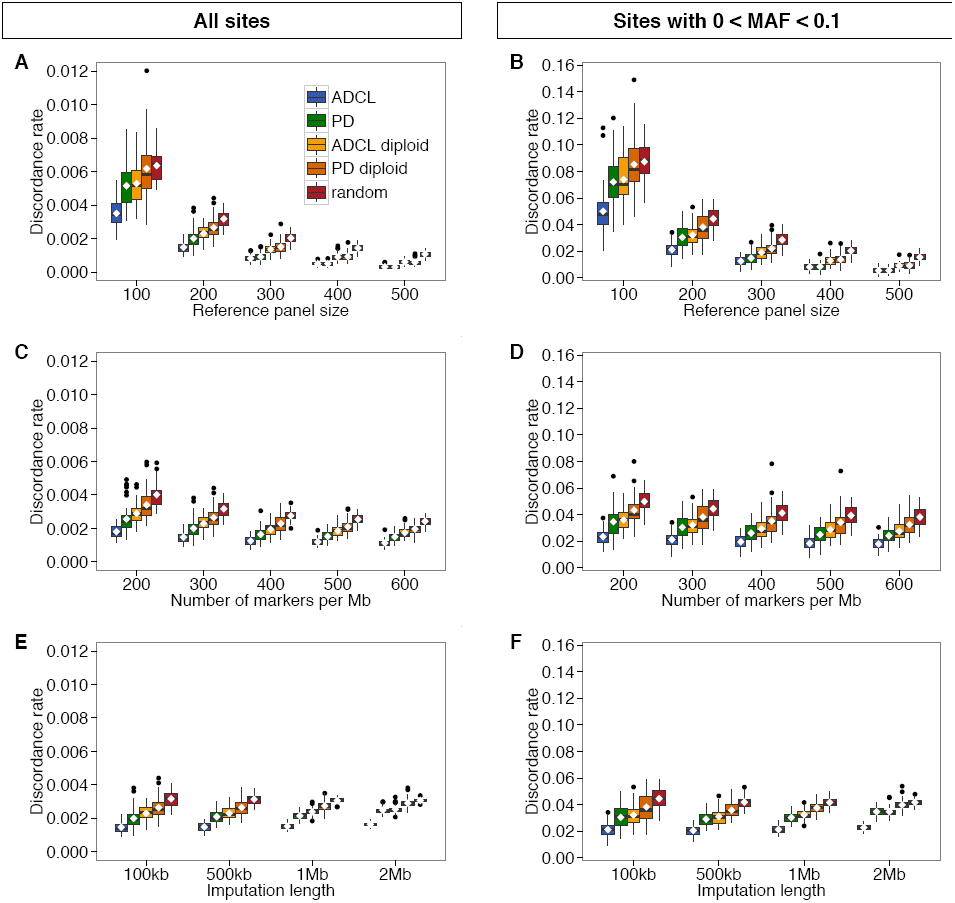
Box plots of discordance rates between imputed and simulated genotypes using the five different reference panel types. **(A)** Varying reference panel size, all sites. **(B)** Varying reference panel size, heterozygous sites with 0 < MAF < 0.1. **(C)** Varying marker density, all sites. **(D)** Varying marker density, heterozygous sites with 0 < MAF < 0.1. **(E)** Varying imputation length, all sites. **(F)** Varying imputation length, heterozygous sites with 0 < MAF < 0.1.

We first evaluated the influence of reference panel size on imputation accuracy, considering cases with *k* equal to 100, 300, 400, and 500 (compared to the initial analysis with *k* = 200). We observe that as the panel size *k* increases, discordance rates decrease across all reference panel types. However, we also note a decrease in the difference in performance between the ADCL and PD algorithms, in both the haploid (“maximum-PD” and “minimum-ADCL”) and diploid cases. In other words, the gain in imputation accuracy obtained by minimizing ADCL instead of maximizing PD diminishes with large reference panel sizes.

Next, we examined how the initial genotyping density of the markers affected imputation accuracy by considering instances with *s* equal to 200, 400, 500, and 600 (compared to the initial choice of *s* = 300). Here, across all reference panel types, the discordance rates decrease slightly with increasing marker density *s*. Nevertheless, for all densities, both the haploid and diploid ADCL panels consistently outperform their PD counterparts in terms of imputation accuracy across all sites, as well as across only the low-frequency variants.

Finally, we considered whether the length of the target imputation region has an effect on imputation accuracy. We imputed segments of length 500kb, 1Mb and 2Mb (compared to the initial imputation length choice of 100kb). In all cases, a flanking 450kb region was added to each end of the sequence in order to avoid edge effects. We observe that discordance rates remain relatively constant across different imputation lengths. Again, the ADCL panels produce significantly lower discordance rates compared to the PD panels, regardless of the specific choice of imputation length.

### Discordance rates with 1000 Genomes Project sequence data

To confirm that our findings on the simulated data set are also observed when using actual sequence data, we performed a similar analysis for 30 1Mb segments generated on chromosome 20, using 381 diploid individuals with European ancestry from the 1000 Genomes Project. We are again interested in comparing the difference in imputation accuracy achieved by the minimum-ADCL and maximum-PD panels, using the mean discordance rate over 1000 randomly-selected reference panels as a baseline for comparison. The discordance rates appear in FIGURE 7, and their mean values are summarized in TABLE 5. For the three different panel types, FIGURE 8 compares the discordance rates examined in each of the 30 segments over all imputed sites, as well as over only the low-frequency variants.

**TABLE 5:** Mean discordance rates between imputed and 1000 Genomes genotypes, using the maximum-PD and minimum-ADCL panels The table is obtained from the data in FIGURES 7 and 8. The comparison is performed over all sites, all heterozygous sites, and heterozygous sites falling into three different MAF groups. Also shown are the *P*-values of the two-tailed Wilcoxon signed-rank tests comparing the discordance rates of the PD and ADCL reference panels.

|  | PD (%) | ADCL (%) | *P*-value |
| --- | --- | --- | --- |
| All | 0.8643 | 0.8087 | $2.367 \times 10^{-3}$ |
| Heterozygotes | 3.9253 | 3.6041 | $9.301 \times 10^{-3}$ |
| MAF (0, 0.1) | 11.3202 | 10.2176 | 0.0234 |
| MAF [0.1, 0.2) | 3.1695 | 2.7525 | 0.0274 |
| MAF [0.2, 0.5] | 1.7302 | 1.5598 | $8.035 \times 10^{-4}$ |

**FIGURE 7:**
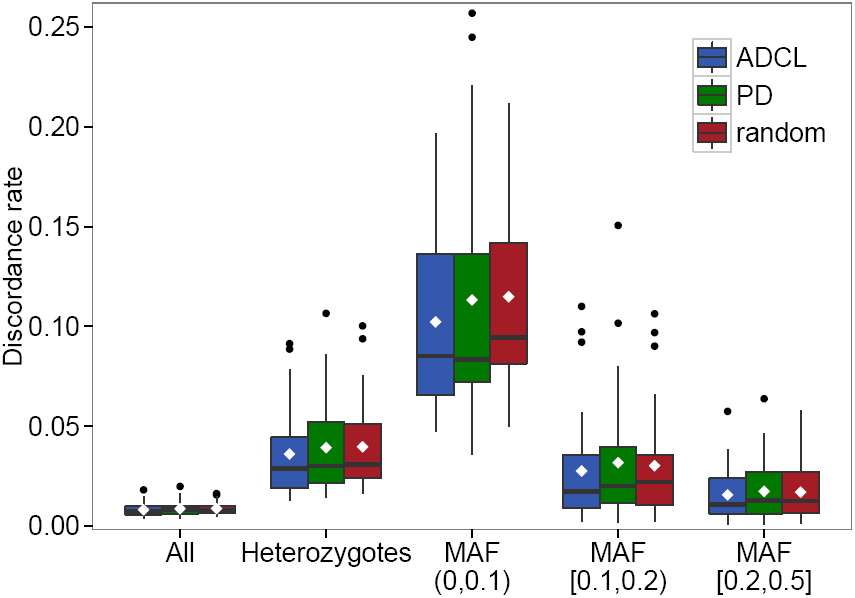
Box plots of discordance rates between imputed and actual genotypes using the minimum-ADCL, maximum-PD, and random panels. The data consist of 30 1Mb segments from 762 haplotypes of European ancestry obtained from the 1000 Genomes Project. The mean discordance rate across the 30 replicates for each comparison group is indicated by a diamond, and the median discordance rate is indicated by a horizontal line. The *x*-axis separates the comparison over all sites, all heterozygous sites, and heterozygous sites falling into three different MAF groups.

**FIGURE 8:**
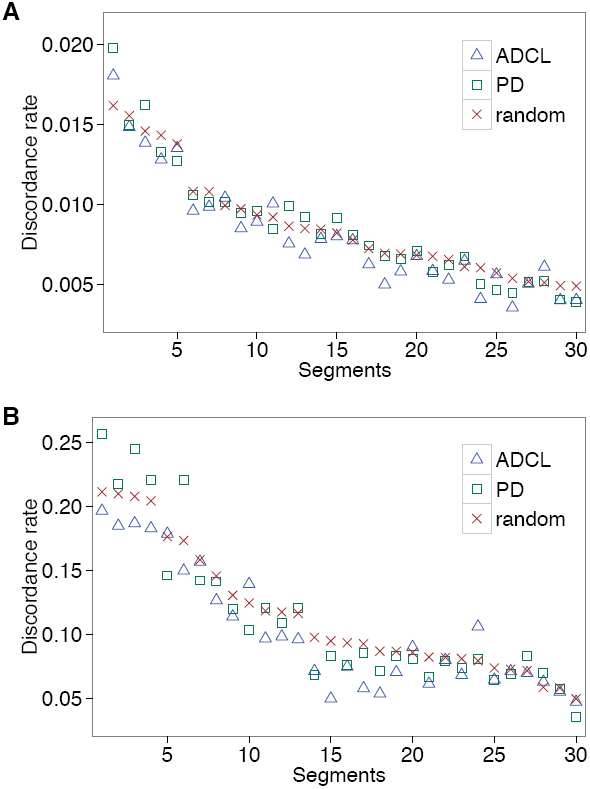
Discordance rates between imputed and actual genotypes using the minimum-ADCL, maximum-PD, and random panels, showing an alternative presentation of the same data used to generate FIGURE 7. **(A)** All sites. **(B)** Heterozygous sites with 0 < MAF < 0.1. The 30 segments are sorted in decreasing order by the mean discordance rate over 1000 random panels.

Applying the two-tailed Wilcoxon signed-rank test, we observe that across all imputed sites, the minimum-ADCL algorithm produces significantly lower discordance rates than the maximum-PD algorithm (*P* = 2.367 × 10^−3^), as shown in TABLE 5. In addition, when focusing solely on the low-frequency variants, the minimum-ADCL panel continues to produce better imputation accuracy than the maximum-PD panel (*P* = 0.0234).

## DISCUSSION

The decreasing cost of modern sequencing has enhanced the practicality of generating a reference panel from the haplotypes that are already present in the study sample. It generally remains prohibitive, however, to perform full sequencing for large numbers of haplotypes. Given this constraint in resources, what is the optimal approach for selecting the subset of the study sample to sequence in order to achieve the best imputation results? We explored two objective functions for optimization, with the aim of ensuring high imputation accuracy.

Maximizing PD as a way of ensuring that the total genetic diversity of a sample is well-represented is one sensible approach. This type of panel selection method achieves lower imputation discordance rates than assembling reference panels from randomly-selected haplotypes (KANG AND MARJORAM 2012; ZHANG *et al*. 2013). Nevertheless, it has not been clear whether PD represents the best objective function for panel selection.

Minimizing ADCL attempts to ensure that the subset of the study sample selected for the panel is representative of the total diversity present, albeit using a different approach. It is conceptually similar to a clustering problem, in that the number of clusters is predetermined, and the algorithm returns the cluster to which each haplotype belongs, as well as the haplotype that is the most central within its cluster. This haplotype is then included in the reference panel. Unlike when maximizing PD, the problem of selecting non-representative branches is mostly avoided by ADCL, as those haplotypes are unlikely to occupy a central position within their clusters.

For both simulated and actual sequence data, we observed that minimizing ADCL does in fact provide an improvement in imputation accuracy compared to maximizing PD. It generally identified a greater number of polymorphic sites, both in the reference panels as well as in the imputed data. When looking at the overall discordance-rate measures, minimizing ADCL produces a significantly lower discordance rate over all sites compared to maximizing PD. This result holds across various choices of genotyping density and imputation length, suggesting that the observed result is robust to such changes. It is only with increasing panel sizes that the gain in imputation accuracy obtained by minimizing ADCL decreases compared to maximizing PD. This outcome could potentially be due to the diminishing returns, in terms of representative variants, contributed by each additional haplotype in the reference panel. Consider the extreme case, where all the haplotypes in the study sample are included in the reference panel. In such a situation, both algorithms return trivially identical imputation results.

One metric that is of particular interest is the performance of an algorithm in the imputation of low-frequency variants. Although early genome-wide association (GWA) studies focused on identifying common variants associated with particular diseases or phenotypic traits, the focus of GWA studies has increasingly shifted toward an interest in rare genetic variants (ASIMIT AND ZEGGINI 2010; CIRULLI AND GOLDSTEIN 2010; EICHLER *et al*. 2010). As such studies improve in their ability to detect the effects of rare variants on phenotype (LI *et al*. 2013; LEE *et al*. 2014), it is paramount that the imputation process carried out alongside them generate reasonably accurate imputed genotypes with low-frequency variants.

In this context, from TABLES 2 and 5, we observed, based on differences in the mean discordance rates, that minimizing ADCL improves upon maximizing PD by the largest absolute amount in the low-MAF bin (0 < MAF < 0.1), in both the simulated and the actual data. This result might be explained by the fact that the discordance rates obtained when imputing low-frequency variants are relatively high to begin with, and can be potentially reduced to a much greater extent with an improved choice of algorithm for panel selection. Our analyses are consistent in suggesting that the minimum-ADCL algorithm can contribute to reducing imputation inaccuracies in GWA studies that seek to identify the effects of low-frequency variants on phenotypic traits.

In summary, we have demonstrated that internal reference panel selection via minimizing ADCL produces empirically improved imputation accuracy compared to maximizing PD, particularly for low-frequency variants. This finding applies to both simulated and actual sequence data, and is robust to changes in the choice of initial parameter values. Note that both ADCL and PD represent intermediate criteria that provide practical objective functions, where the ultimate goal is maximizing imputation accuracy or other aspects of imputation performance. Although both algorithms produce considerably better imputation performance measures than the use of random panels, neither is guaranteed to produce the maximal value of such measures over all possible panels. It remains to be determined whether a single simple criterion exists that could lead to identification of the best possible panel for maximizing imputation performance.

## ACKNOWLEDGMENTS

This work was supported by National Institutes of Health grant R01 HG005855.

## LITERATURE CITED

Asimit J., and E. Zeggini, 2010. Rare variant association analysis methods for complex traits. Annu. Rev. Genet. 44: 293–308.

Bordewich M., A. G. Rodrigo, and C. Semple, 2008. Selecting a taxa to save or sequence: desirable criteria and a greedy solution. Syst. Biol. 57: 825–834.

Cirulli E. T., and D. B. Goldstein, 2010. Uncovering the roles of rare variants in common disease through whole-genome sequencing. Nat. Rev. Genet. 11: 415–425.

Eichler E. E., J. Flint, G. Gibson, A. Kong, S. M. Leal et al., 2010. Missing heritability and strategies for finding the underlying causes of complex disease. Nat. Rev. Genet. 11: 446–450.

Faith D. P., 1992. Conservation evaluation and phylogenetic diversity. Biol. Conserv. 61: 1–10.

Fridley B. L., G. Jenkins, M. E. Deyo-Svendsen, S. Hebbring, and R. Freimuth, 2010. Utilizing genotype imputation for the augmentation of sequence data. PLoS ONE 5: e11018.

Hartmann K., and M. Steel, 2007. Phylogenetic diversity: from combinatorics to ecology, pp. 171–196 in Reconstructing Evolution: New Mathematical and Computational Advances, edited by O. Gascuel, and M. Steel. Oxford University Press, Oxford.

Howie, B., C. Fuchsberger, M. Stephens, J. Marchini, and G. R. Abecasis, 2012. Fast and accurate genotype imputation in genome-wide association studies through pre-phasing. Nat. Genet. 44: 955–959.

Huang, L., Y. Li, A. B. Singleton, J. A. Hardy, G. Abecasis et al., 2009. Genotype-imputation accuracy across worldwide human populations. Am. J. Hum. Genet. 84: 235–250.

Huang, L., M. Jakobsson, T. J. Pemberton, M. Ibrahim, T. Nyambo et al., 2011. Haplotype variation and genotype imputation in African populations. Genet. Epidemiol. 35: 766–780.

Hudson, R. R., 2002. Generating samples under a Wright-Fisher neutral model of genetic variation. Bioinformatics 18: 337–338.

International HapMap Consortium, 2005. A haplotype map of the human genome. Nature 437: 1299–1320.

Jewett, E. M., M. Zawistowski, N. A. Rosenberg, and S. Zöllner, 2012. A coalescent model for genotype imputation. Genetics 191: 1239–1255.

Kang, C. J., and P. Marjoram, 2012. A sample selection strategy for next-generation sequencing. Genet. Epidemiol. 36: 696–709.

Kaufman, L., and P. J. Rousseeuw, 1987. Clustering by means of medoids, pp. 405–416 in Statistical Data Analysis Based on the L1-Norm and Related Methods, edited by Y. Dodge. North-Holland, Amsterdam.

Kreiner-Møller, E., C. Medina-Gomez, A. G. Uitterlinden, F. Rivadeneira, and K. Estrada, 2015. Improving accuracy of rare variant imputation with a two-step imputation approach. Eur. J. Hum. Genet. 23: 395–400.

Lee, S., G. R. Abecasis, M. Boehnke, and X. Lin, 2014. Rare-variant association analysis: study designs and statistical tests. Am. J. Hum. Genet. 95: 5–23.

Li, B., D. J. Liu, and S. M. Leal, 2013. Identifying rare variants associated with complex traits via sequencing. Curr. Protoc. Hum. Genet. 78: 1.26.1–1.26.22.

Li, Y., C. Willer, S. Sanna, and G. Abecasis, 2009. Genotype imputation. Annu. Rev. Genomics Hum. Genet. 10: 387–406.

Li, Y., C. J. Willer, J. Ding, P. Scheet, and G. R. Abecasis, 2010. MaCH: using sequence and genotype data to estimate haplotypes and unobserved genotypes. Genet. Epidemiol. 34: 816–834.

Marchini, J., and B. Howie, 2010. Genotype imputation for genome-wide association studies. Nat. Rev. Genet. 11: 499–511.

Matsen, F. A., A. Gallagher, and C. O. McCoy, 2013. Minimizing the average distance to a closest leaf in a phylogenetic tree. Syst. Biol. 62: 824–836.

Pardi, F., and N. Goldman, 2005. Species choice for comparative genomics: being greedy works. PLoS Genet. 1: e71.

Paşaniuc, B., R. Avinery, T. Gur, C. F. Skibola, P. M. Bracci et al., 2010. A generic coalescent-based framework for the selection of a reference panel for imputation. Genet. Epidemiol. 34: 773–782.

Peil, B., M. Kabisch, C. Fischer, U. Hamann, and J. L. Bermejo, 2015. Tailored selection of study individuals to be sequenced in order to improve the accuracy of genotype imputation. Genet. Epidemiol. 39: 114–121.

Saitou, N., and M. Nei, 1987. The neighbor-joining method: a new method for reconstructing phylogenetic tress. Mol. Biol. Evol. 4: 406–425.

Sampson J. N., K. Jacobs, Z. Wang, M. Yeager, S. Chanock et al., 2012. A two-platform design for next generation genome-wide association studies. Genet. Epidemiol. 36: 400–408.

Sheng, W., and X. Liu, 2004. A hybrid algorithm for *k*-medoid clustering of large data sets. Proc. IEEE Congr. Evol. Comput. 1: 77–82.

Shriner, D., and A. Adeyemo, G. Chen, C. N. Rotimi, 2010. Practical considerations for imputation of untyped markers in admixed populations. Genet. Epidemiol. 34: 258–265.

Simonsen, M., T. Mailund, and C. N. S. Pedersen, 2008. Rapid neighbor-joining, pp. 113–122 in Algorithms in Bioinformatics, edited by K. A. Crandall, and J. Lagergren. Springer-Verlag, Berlin.

Steel, M., 2005. Phylogenetic diversity and the greedy algorithm. Syst. Biol. 54: 527–529.

Sukumaran, J., and M. T. Holder, 2010. DendroPy: A Python library for phylogenetic computing. Bioinformatics 26: 1569–1571.

Surakka, I., K. Kristiansson, V. Anttila, M. Inouye, C. Barnes et al., 2010. Founder population-specific HapMap panel increases power in GWA studies through improved imputation accuracy and CNV tagging. Genome Res. 20: 1344–1351.

The 1000 Genomes Project Consortium, 2010. A map of human genome variation from population-scale sequencing. Nature 467: 1061–1073.

Theodoridis, S., and K. Koutroumbas, 2008. Pattern Recognition (4th ed.). Academic Press, Waltham, MA.

Zhang, P., X. Zhan, N. A. Rosenberg, and S. Zöllner, 2013. Genotype imputation reference panel selection using maximal phylogenetic diversity. Genetics 195: 319–330.

